# Effects of Healthy Aging on Right Ventricular Structure and Biomechanical Properties

**DOI:** 10.1101/2020.09.08.288332

**Authors:** Danial Sharifi Kia, Yuanjun Shen, Timothy N. Bachman, Elena A. Goncharova, Kang Kim, Marc A. Simon

**Author notes:** **Address correspondence to:** Marc A. Simon, MD, MSc, Associate Professor of Medicine, Division of Cardiology, Department of Medicine, University of Pittsburgh School of Medicine, Presbyterian University Hospital C-701, 200 Lothrop St, Pittsburgh, PA.

## Abstract

Healthy aging has been associated with alterations in pulmonary vasculature and right ventricular (RV) hemodynamics, potentially leading to RV remodeling. Despite the current evidence suggesting an association between aging and alterations in RV function and higher prevalence of pulmonary hypertension in the elderly, limited data exist on age-related differences in RV structure and biomechanics. In this work we report our preliminary findings on the effects of healthy aging on RV structure, function, and biomechanical properties. Hemodynamic measurements, biaxial mechanical testing, constitutive modeling, and quantitative histological analysis were employed to study two groups of Sprague-Dawley rats: control (11 weeks) and aging (80 weeks).

Aging was associated with increases in RV peak pressures (≈↑17%, p=0.017), RV contractility (≈↑52%, p= 0.004), and RV wall thickness (≈↑34%, p=0.002). Longitudinal realignment of RV collagen (16.4°, p=0.013) and myofibers (14.6°, p=0.017) were observed with aging, accompanied by transmural cardiomyocyte loss and fibrosis. A bimodal alteration in biomechanical properties was noted, resulting in increased myofiber stiffness (≈↑158%, p=0.0006) and decreased effective collagen fiber stiffness (≈↓67%, p=0.031).

Our results demonstrate the potential of healthy aging to modulate RV remodeling via increased peak pressures, cardiomyocyte loss, fiber reorientation, and altered collagen/myofiber stiffness. Some similarities were observed between aging-induced remodeling patterns and those of RV remodeling in pressure overload.

## 1. Introduction

Healthy aging is associated with alterations in right ventricular (RV) structure and function in subjects with no underlying cardiopulmonary disease (D’Andrea et al., 2017; Fiechter et al., 2013; Granath et al., 1964; Innelli et al., 2009; Nakou et al., 2016). Aging results in pulmonary artery (PA) remodeling (Hosoda et al., 1984; Sicard et al., 2018), increased pulmonary vascular resistance (Ehrsam et al., 1983; Granath et al., 1964) and increased PA systolic pressures (Kane et al., 2016; Lam et al., 2009), leading to RV remodeling (Anversa et al., 1990; Chouabe et al., 2004), altered contraction dynamics (Effron et al., 1987), and decreased RV systolic strains (Chia et al., 2014). Moreover, aging results in diminished RV hypertrophy in response to pressure overload (Chouabe et al., 2004; Kuroha et al., 1991). Age-related differences exist in the 1, 2 and 3-year survival rates of pulmonary hypertension (PH) patients (Hoeper et al., 2013).

In recent years, biomechanical analysis techniques have been employed to better understand the underlying mechanisms of RV remodeling (Avazmohammadi et al., 2017b, 2019; Hill et al., 2014; Sharifi Kia et al., 2020) and have closely linked RV biomechanics to function (Jang et al., 2017). Despite the evidence suggesting an association between aging and alterations in RV structure/function and higher prevalence of PH in the elderly (Hoeper and Gibbs, 2014), a large body of the literature has employed younger animal models and limited data exist on age-associated differences in RV biomechanics.

In this short communication we present our preliminary findings on the effects of healthy aging on RV structure, function, and biomechanical properties. Our study provides insights into how healthy aging modulates RV remodeling and lays the ground for future work to study the age-associated differences in RV response to pressure overload.

## 2. Methods

The data acquired during this study are available from the corresponding author on reasonable request. A total of 15 male Sprague-Dawley rats corresponding to young (controls, ≈11 weeks, *n_Control_* = 9) and old (≈80 weeks, *n_Aging_* = 6) age groups were studied using a multi-scale biomechanical analysis framework. **Figure 1** summarizes the experimental procedures and different analysis techniques used in this work. All animal procedures were approved by University of Pittsburgh’s IACUC (protocol# 18113872 and 19126652). Details of our experimentation/analysis techniques have been extensively discussed in prior studies (Hill et al., 2014; Sharifi Kia et al., 2020). Please see supplementary materials for details on our methods.

**Figure 1:**
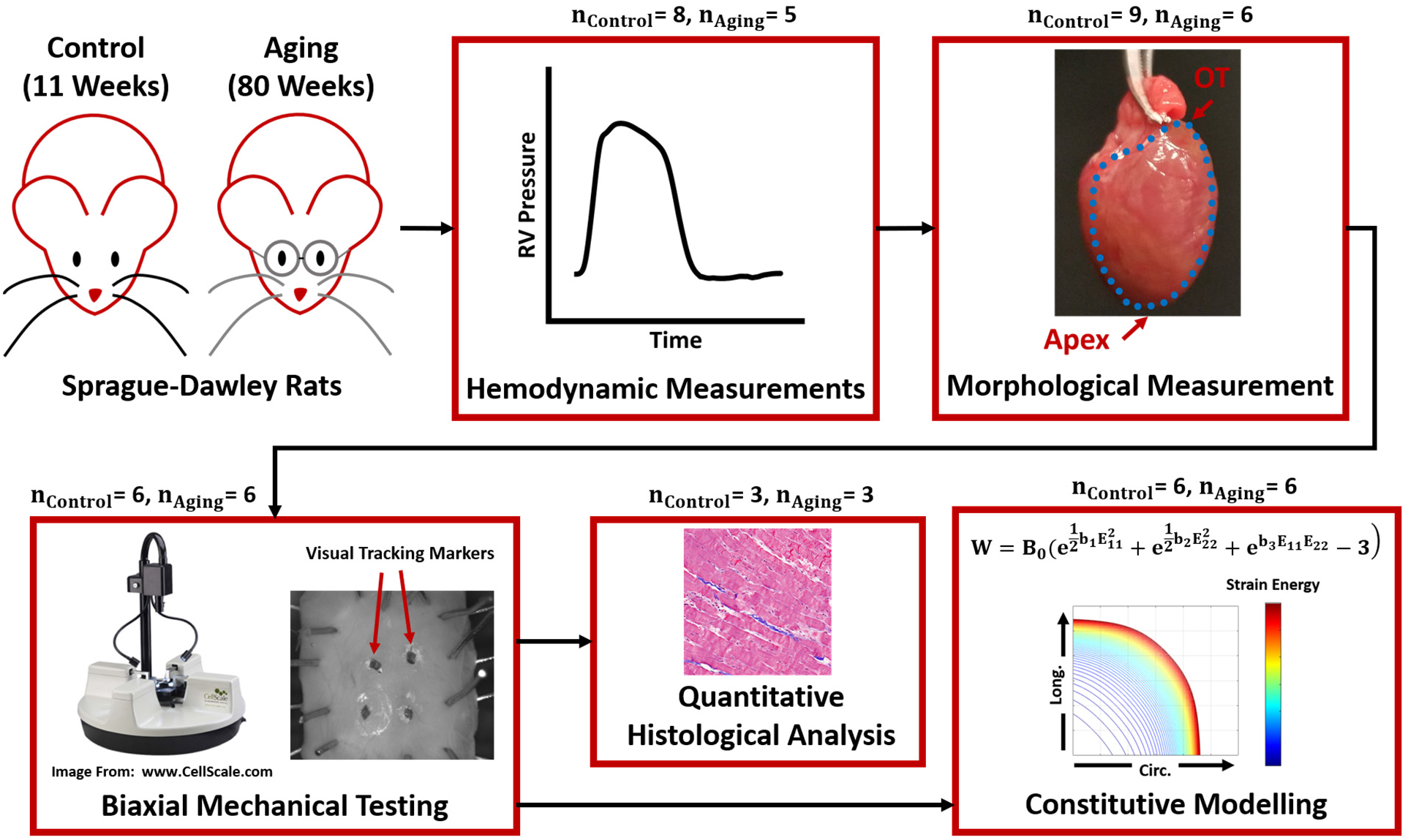
The framework used to study RV remodeling with healthy aging. *In vivo* terminal invasive hemodynamic measurements were performed on young controls and aging Sprague-Dawley rats, followed by harvesting the heart and morphological measurements, biaxial mechanical testing, constitutive modeling, and quantitative transmural histological analysis. RV: Right ventricle; OT: Outflow tract; Circ: Circumferential; Long.: Longitudinal

Briefly, terminal invasive pressure catheterization was performed on both groups and analyzed for common metrics of RV hemodynamics. The hearts were then harvested, and measurements were acquired for the Fulton index and RV free wall (RVFW) thickness.

To characterize the effects of aging on RV biomechanical properties, multi-protocol biaxial mechanical testing was performed on RVFW specimens, followed by estimation of effective fiber-ensemble biomechanical properties of myofibers and collagen. Additionally, a nonlinear anisotropic constitutive model (Choi and Vito, 1990) was used for modeling the response of each specimen to generate age-specific strain energy maps of RVFW mechanical properties.

Transmural histological staining (Masson’s trichrome) was performed on a sub-group of specimens, to quantify the effects of aging on RV fiber architecture (collagen stained in blue, myofibers strained in red/pink). Cardiomyocyte width was then measured from the histological sections for each group. Additionally, transmural area fractions of collagen/myofibers were quantified for each section to analyze the effects of aging on RVFW composition. Furthermore, transmural orientations of collagen and myofibers were quantified using gradient-based techniques. Collagen fiber coherency was measured transmurally to analyze the effects of aging on collagen morphology.

Data are presented with mean ± standard error of the mean. Sample normality and homogeneity of variance were assessed using the Shapiro–Wilk test and Bartlett’s test of homoscedasticity. Circular statistics was employed for analysis of fiber orientation data, using the Watson–Williams test in CircStat toolbox (Berens, 2009) in MATLAB (Mathworks, Natick, MA). For all other data, in case of normality and homoscedasticity, a two-sided unpaired student’s t-test was used for statistical comparisons. Welch’s t-test was used for non-homoscedastic data, while non-normal distributions were compared using Mann–Whitney U-tests. For all purposes, p<0.05 was considered statistically significant. Statistical analyses were performed in the R software package (R Foundation for Statistical Computing, Vienna, Austria).

## 3. Results

### 3.1. RV Hemodynamics and Morphology

Healthy aging did not show an effect on heart rate (**Fig. 2a**; 271.5±11.7 vs. 292.3±14.1 BPM for Aging-vs.-Control; p=0.326). Aging resulted in increased RV peak pressures (**Fig. 2b**; 26.8±0.9 vs. 23.0±0.9 mmHg for Aging-vs.-Control; p=0.017), while showing a modest non-significant effect on end-diastolic pressures (**Fig. 2c**; 1.9±0.4 vs. 1.3±0.1 mmHg for Aging-vs.-Control; p=0.085). Effects of aging on load-dependent measures of RV contractility and relaxation are shown in **Figure 2d**. Aging significantly increased 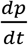max (1611.7±90.5 vs. 1063.8±101.7 mmHg/s for Aging-vs.-Control; p=0.004), but did not demonstrate any effects on 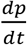min (−823.9±60.4 vs. −814.7±85.5 mmHg/s for Aging-vs.-Control; p=0.940). Increased contractility index was observed for the aging group (**Fig. 2e**; 60.1±2.2 vs. 45.8±3.5 1/s for Aging-vs.-Control; p=0.012), while the time constant of RV relaxation (tau) remained unchanged (**Fig. 2f**; 10.7±1.6 vs. 9.9±0.8 ms for Aging-vs.-Control; p=0.595). As shown in **Table 1**, healthy aging significantly increased RV wall thickness (0.89±0.06 vs. 0.66±0.03 mm for Aging-vs.-Control; p=0.002), while not affecting the Fulton index (0.26±0.03 vs. 0.27±0.01 mg/mg for Aging-vs.-Control; p=0.140).

**Figure 2:**
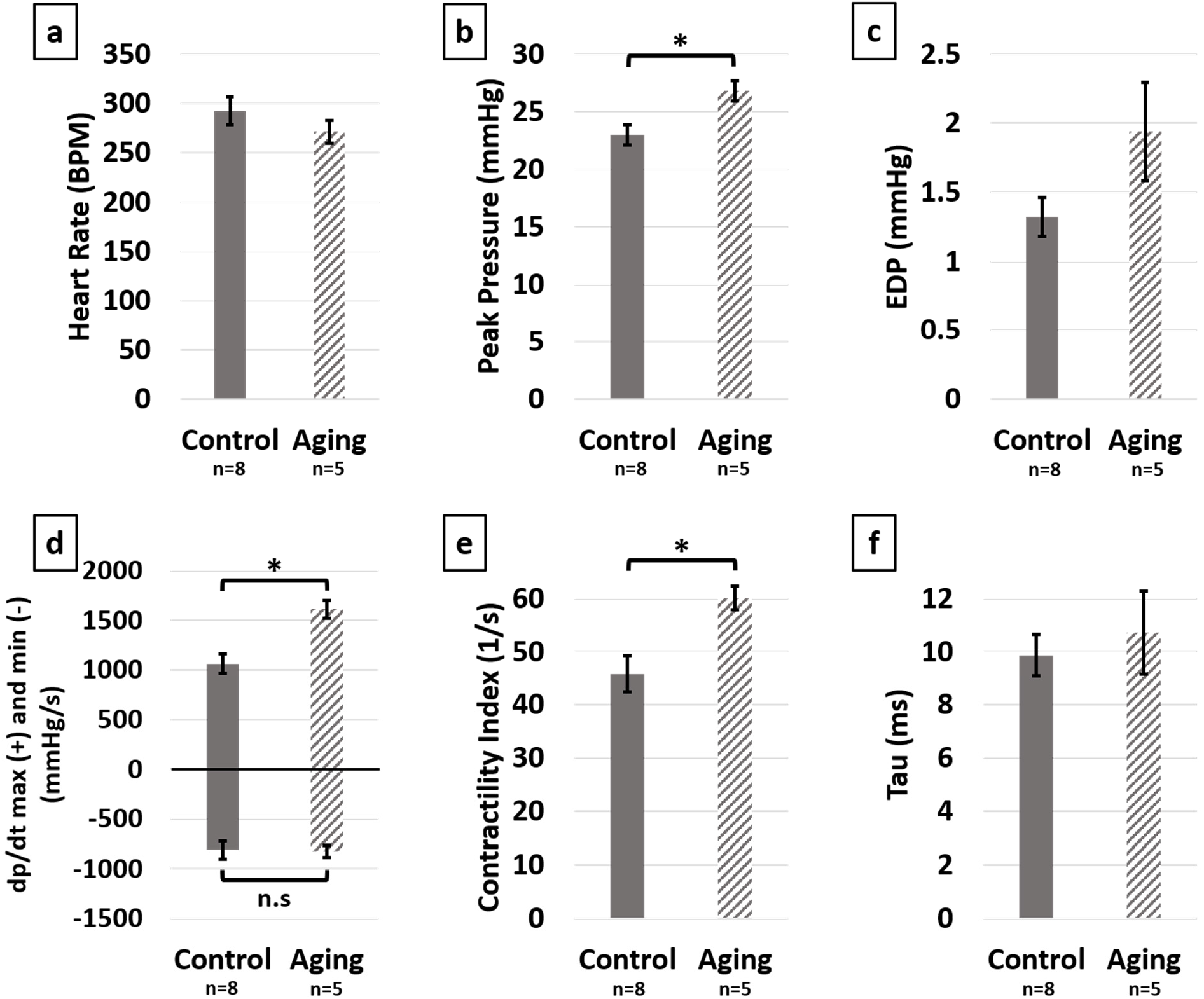
Hemodynamic measures of the effects of healthy aging on RV **(a)** Heart rate, **(b)** Peak pressure, dp **(c)** End-diastolic pressure, **(d)** 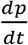 (positive side) and 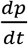 (negative side), **(e)** Contractility index and **(f)** Preload-independent measure of relaxation (tau). Healthy aging significantly increased RV peak pressures and the load-dependent measures of RV contractility (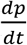 and contractility index), while not affecting the heart rate, end-diastolic pressures and relaxation function (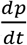 and tau). Error bars represent standard error of the mean (SEM). * indicates p<0.05. RV: Right ventricle; BPM: Beats per minute; EDP: End-diastolic pressure; 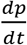 and min: Load-dependent measures of RV contractility and relaxation; n.s: Non-significant.

**Table 1:** Effects of healthy aging on RV hypertrophy (RVFW thickness) and Fulton index

|  | **RVFW Thickness**<br>**(mm)** | **Fulton Index**<br>**(mg/mg)** |
| --- | --- | --- |
| **Control**<br>**(n=9)** | 0.66±0.03 | 0.27±0.01 |
| **Aging**<br>**(n=6)** | 0.89±0.06 | 0.26±0.03 |
| p Value | 0.002 | 0.140 |

### 3.2. RVFW Biomechanical Properties

Aging demonstrated a bimodal effect on RVFW biaxial properties by resulting in increased circumferential and longitudinal stiffness under lower strains, while progressing to decreased biaxial stiffness at higher strains (**Fig. 3a**). A similar effect was observed on the effective fiber-ensemble stress-strain properties of RVFW collagen and myofibers (**Fig. 3b**). Using a rule of mixtures approach, this translated into increased effective myofiber stiffness (**Fig. 3c**; 169.9±20.6 vs. 65.8±4.7 kPa for Aging-vs.-Control; p=0.0006) and decreased effective collagen fiber stiffness (**Fig. 3d**; 23.9±3.8 vs. 73.1±15.4 MPa for Aging-vs.-Control; p=0.031). Constitutive model parameters for each group are shown in **Table 2**. Overall, the employed model showed an acceptable fit quality to our experimental data (R^2^=0.95±0.01 and 0.96±0.01 for Aging and Control, respectively). Age-specific strain energy maps, representing the combined effects of all model parameters, are demonstrated in **Figure 3e-f** for the low-strain and high-strain regions.

**Figure 3:**
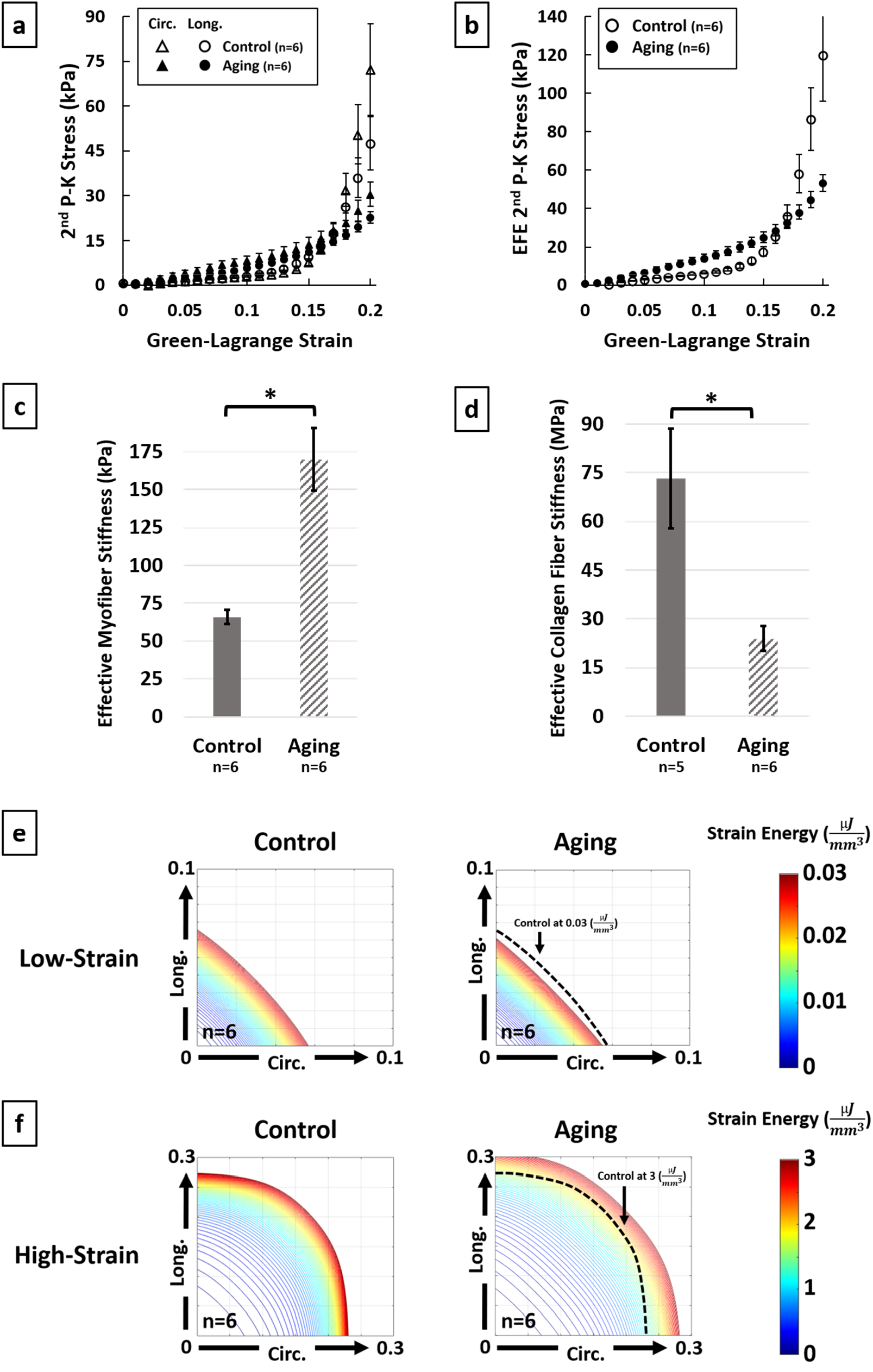
Effects of healthy aging on **(a)** Biaxial mechanical properties of RV myocardium, **(b)** Effective fiber-ensemble mechanical properties of combined myofiber–collagen bundles, **(c)** Effective myofiber stiffness, **(d)** Effective collagen fiber stiffness, **(e)** Strain energy maps of RVFW in the low-strain region (circumferential-longitudinal strain space) and **(f)** Strain energy maps of RVFW in the high-strain region. Healthy aging modulates the biomechanical properties of RVFW in a bimodal manner by increasing the effective myofiber stiffness while decreasing collagen fiber stiffness. Strain energy maps demonstrate increased RVFW stiffness in the low-strain region with healthy aging (mostly dominated by myofibers), followed by decreased RVFW stiffness in the high-strain region post-collagen recruitment. Error bars represent standard error of the mean (SEM).* indicates p<0.05. RV: Right ventricle; RVFW: Right ventricular free wall; 2^nd^ P-K Stress: 2^nd^ Piola-Kirchhoff stress; Circ.: Circumferential; Long.: Longitudinal; EFE 2^nd^ P-K Stress: Effective fiber-ensemble 2^nd^ Piola-Kirchhoff stress.

**Table 2:** Constitutive model parameters representing the circumferential (b1), longitudinal (b_2_) and in-plane coupling (b_3_) stiffness of RV myocardium for the control and aging groups

|  | **Model Parameters** |  |  |  |
| --- | --- | --- | --- | --- |
|  | **B<sub>0</sub>** | **b<sub>1</sub> (kPa)** | **b<sub>2</sub> (kPa)** | **b<sub>3</sub> (kPa)** |
| **Control**<br>**(n=6)** | 0.173±0.023 | 98.23±14.52 | 76.33±7.53 | 66.72±6.64 |
| **Aging**<br>**(n=6)** | 0.369±0.082 | 56.01±7.12 | 47.35±6.88 | 43.21±3.80 |
| p Value | 0.044 | 0.026 | 0.017 | 0.012 |

### 3.3. Quantitative Transmural Histology

Representative histological sections are demonstrated for each group in **Figure 4a**. Aging resulted in increased cardiomyocyte width (**Fig. 4b**; 25.42±0.34 vs. 14.94±0.64 μm for Aging-vs.-Control; p=0.0001). Quantifying the transmural orientation of RVFW fibers revealed myofiber (**Fig. 4c**) and collagen (**Fig. 4d**) reorientation towards the longitudinal direction at sub-endocardial levels. Overall, myofibers showed similar orientations to collagen fibers. Aging significantly shifted the overall orientation of myofibers (circular mean of transmural distributions, dotted lines in **Fig. 4c**) by 14.6° towards the longitudinal direction (**Fig. 4e**; p=0.017). Similarly, the dominant orientation of collagen fibers was shifted by 16.4° (p=0.013). Aging also resulted in cardiomyocyte loss and decreased myofiber area fractions at both epicardium (**Fig. 4f**; 90.8±0.3% vs. 95.3±0.7% for Aging-vs.-Control; p=0.004) and endocardium (**Fig. 4f**; 82.4±1.5% vs. 95.3±1.9% for Aging-vs.-Control; p=0.007), with a non-significant effect on the mid-ventricular region (**Fig. 4f**; 91.8±1.3% vs. 95.4±0.5% for Aging-vs.-Control; p=0.063). Furthermore, aging lead to RVFW fibrosis and increased collagen area fractions at epicardium (**Fig. 4g**; 5.3±0.4% vs. 3.4±0.3% for Aging-vs.-Control; p=0.015) and the mid-ventricular region (**Fig. 4g**; 5.0±0.4% vs. 3.4±0.3% for Aging-vs.-Control; p=0.037). Analyzing the coherency of collagen architectures revealed decreased coherency at the endocardium (**Fig. 4h**; 10.4±1.1% vs. 19.7±1.1% for Aging-vs.-Control; p=0.003), while not showing any effects on other transmural regions.

**Figure 4:**
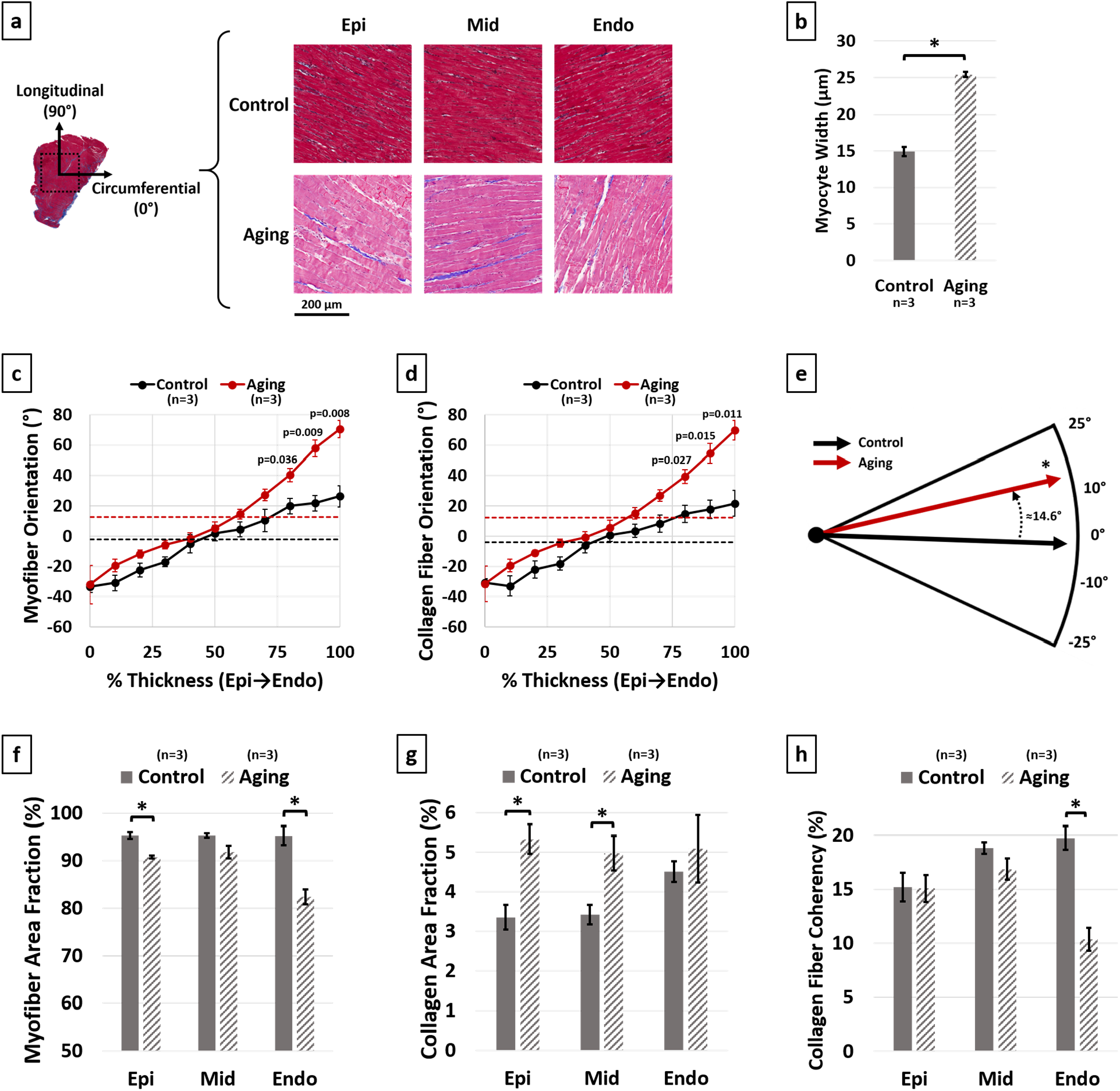
Histological analysis of the effects of healthy aging on RV structure. **(a)** Representative transmural histological sections of the RVFW (Red/Pink: Myofiber, Blue: Collagen) and effects of aging on **(b)** Cardiomyocyte hypertrophy (myocyte width), **(c)** Transmural myofiber orientations, **(d)** Transmural collagen fiber orientations, **(e)** Dominant myofiber orientations, **(f)** Transmural myofiber content (area fraction), **(g)** Transmural collagen content (area fraction) and **(h)** Transmural collagen fiber coherency. Healthy aging results in cardiomyocyte hypertrophy, in addition to reorientation of sub-endocardial collagen and myofibers towards the longitudinal direction. This is accompanied by loss of cardiomyocytes, RVFW fibrosis and decreased collagen fiber coherency at endocardium. Error bars represent standard error of the mean (SEM). * indicates p<0.05. RV: Right ventricle; RVFW: Right ventricular free wall; Epi: Epicardium; Mid: Mid-ventricular region; Endo: Endocardium.

## 4. Discussion

We aimed to investigate the effects of healthy aging on RV remodeling. Aging resulted in **1)** Increased peak pressures (≈↑17%) and RV contractility (≈↑52%). **2)** Increased RVFW thickness (≈↑34%) with no effects on the Fulton index, indicating proportional right and left ventricular growth. **3)** Longitudinal reorientation of collagen/myofibers, together with cardiomyocyte loss and RVFW fibrosis with decreased collagen fiber coherency. **4)** Increased effective myofiber stiffness (≈↑158%) accompanied by decreased collagen fiber stiffness (≈↓67%).

The observed increase in RV peak pressures (**Fig. 2b**) are consistent with previous reports of increased PA systolic pressures and RV afterload with healthy aging (Kane et al., 2016; Lam et al., 2009). Furthermore, cardiomyocyte width (**Fig. 4b**) and RVFW thickness (**Table 1**) increased with aging, leading to increased organ-level contractility (**Fig. 2d-e**). This is similar to previous observations of RV hypertrophy in PH and increased contractility to overcome the elevated afterload (Hill et al., 2014; Sharifi Kia et al., 2020).

Histological analyses revealed reorientation of endocardial collagen and myofibers, resulting in a longitudinal shift in dominant transmural orientation of RVFW fibers (**Fig. 4e**). Similar patterns of fiber reorientation were noted in animal models of PH (Avazmohammadi et al., 2017a; Sharifi Kia et al., 2020). Consistent with prior work (Anversa et al., 1990; Walker et al., 2006), cardiomyocyte loss (**Fig. 4f**) and RVFW fibrosis (**Fig. 4g**) were noted due to aging. Cardiomyocyte loss in the RVFW increases the hemodynamic load on the remaining myocytes (Fajemiroye et al., 2018) and may explain the observed hypertrophy patterns (**Fig. 4b**). While requiring further experimentation, a potential explanation for RV myocyte loss and extracellular matrix (ECM) deposition may be local RVFW ischemia due to aging (Anversa et al., 1990).

Fiber reorientation (**Fig. 4 c-d**) and increased transmural change in fiber angles, lead to a less anisotropic biaxial mechanical response for the aging group (**Fig. 3a**). Bimodal alterations in RVFW biaxial properties were observed with aging (**Fig. 3a**). Compared to controls, the aging group showed higher stiffness in the low-strain region, followed by lower biaxial stiffness at higher strains. Moreover, aging lead to increased effective myofiber stiffness (**Fig. 3c**). Increased myofiber stiffness and reduced tissue-level ventricular stiffness at high strains have been previously documented in separate studies on age-related left ventricular remodeling (Cappelli et al., 1984; Lieber et al., 2004). Potential underlying mechanisms of increased myofiber stiffness include myocyte remodeling due to cell loss, as well as reduced titin phosphorylation (Rain et al., 2013). Despite an increase in collagen area fractions (**Fig. 4g**), effective collagen fiber stiffness decreased with aging (**Fig. 3d**). In this study, collagen stiffness measures represent the effective response of collagen fibers, estimated from the tissue-level behavior. Therefore, while this could indicate a decrease in the intrinsic stiffness of collagen fibers, it does not eliminate the possibility of tissue-level structure/architecture affecting the observed behavior (Hill et al., 2014). For instance, reduced collagen fiber coherency was detected at the endocardial levels (**Fig 4h**), indicating a more sparse and isotropic distribution of collagen fibers in the aging group (Clemons et al., 2018). This has the potential to affect the load transfer mechanism of endocardial collagen fibers, contributing to reduced effective modulus at the tissue level. Further experimentation is warranted on the mechanisms of collagen fiber remodeling with aging. Candidate mechanisms for decreased effective collagen stiffness include 1) Lysyl oxidase- mediated alterations in collagen cross-linking, 2) Reduced effective collagen stiffness due to myocyte loss and altered biomechanics in the collagen/myofiber niche, hampering effective load transfer in the RVFW continuum, 3) Altered collagen microstructure and crimp, resulting in delayed collagen recruitment.

Age-specific strain energy maps were employed to investigate the effects of healthy aging on RV biomechanical properties. Model predictions demonstrate a similar bimodal strain energy response to experimentally measured data, indicating increased stiffness in the low-strain region, followed by decreased stiffness at higher strains (**Fig. 3e-f**).

Despite low variability and strong statistics, the small sample size of our preliminary study remains a limitation. Moreover, lack of volumetric hemodynamic data prevents analyzing the effects of aging on *in vivo* RV structure. Additionally, we employed a phenomenological constitutive model for analyzing our biomechanical data. Future work will focus on structurally-informed constitutive models of RV myocardium (Avazmohammadi et al., 2017b) to couple the histologically measured tissue architecture to biaxial properties. Different batches of staining solution used for each group, resulted in different shades of cardiomyocyte staining for control vs. aging (red vs. pink). However, this will have minimal effects on our findings since segmentation thresholds for myofibers and collagen were individually selected for each histological section.

In summary, our results demonstrate the potential of healthy aging to modulate RV remodeling via increased peak pressures, cardiomyocyte loss, fibrosis, fiber reorientation and altered collagen/myofiber stiffness. Some similarities were observed between aging-induced remodeling patterns and RV remodeling in PH.

## Acknowledgments

This study was supported by the American Heart Association (AHA-20PRE35210429, D.S.K) and the National Institutes of Health (NIH Grants 1R01AG058659, 2P01HL103455, and UL1 TR001857, M.A.S; 2R01HL130261, 2R01HL113178, and R01HL150638, E.A.G). The funding sources had no involvement in design of the study, data acquisition or interpretation.

