## Supplementary Materials for "Effects of Healthy Aging on Right Ventricular Structure and Biomechanical Properties"

### 1. Supplementary Methods

A total of 15 male Sprague-Dawley rats corresponding to young (≈11 weeks, $n_{Control} = 9$) and old (≈80 weeks, $n_{Aging} = 6$) age groups were studied in this work. An approximately 70 weeks age difference between the control and aging groups was considered sufficient to study the effects of healthy aging on RV structure/function in the absence of pathological events arising with senescence in older animals. Historical data from a recent study in our laboratory (Sharifi Kia et al., 2020) was used for the control animals in this work.

#### 1.1. Hemodynamic and Morphological Measurements

Using standard techniques, terminal invasive pressure catheterization was performed on both groups, using a Millar™ conductance catheter (Millar Inc., Houston, TX) for the young controls ($n_{Control} = 8$) and a Transonic™ conductance catheter (Transonic Systems Inc., Ithaca, NY) for the aging group ($n_{Aging} = 5$). Open-chest hemodynamic measurements were performed under anesthesia induced via inhalation of isoflurane, while the animals were placed on a heated table (37°C) and monitored using a rectal probe. Pressure data was then analyzed for common RV hemodynamic metrics, including heart rate, peak pressures and measures of contractility and relaxation. Following hemodynamic measurements, the heart was harvested and measurements were acquired for the Fulton index (ratio of RV weight to weight of the left ventricle + intraventricular septum) and RV free wall (RVFW) thickness ($n_{Control} = 9$, $n_{Aging} = 6$).

#### 1.2. Biomechanical Characterization

In order to characterize the effects of aging on RV biomechanical properties, following morphological measurements, square specimens were harvested from the RVFW to undergo biaxial testing ($n_{Control} = 6$, $n_{Aging} = 6$). Specimens were mounted on a BioTester biaxial testing device (CellScale, Waterloo, ON, Canada) in trampoline fashion (using a suture and pulley mechanism) to ensure minimal shear loading (Sacks, 1999). Samples were submerged in modified Krebs solution with 2,3-Butanedione monoxime and oxygen to ensure tissue viability (Valdez-Jasso et al., 2012) and underwent multi-protocol biaxial mechanical testing (1:1, 1:2, 2:1, 1:4, 4:1, 1:6 and 6:1 displacement ratios) to investigate the RVFW response in a wide range of possible loading scenarios. Visual tracking markers were placed on the epicardial surface of the RVFW specimens and marker displacements (recorded using a CCD camera) were analyzed to obtain the deformation gradient, Green–Lagrange strain and the 2nd Piola–Kirchhoff stress tensors, using a finite deformation analysis framework in Mathcad (PTC, Needham, MA). Equibiaxial response of the specimens were interpolated from the multi-protocol test data, using well-established techniques (Avazmohammadi et al., 2017b; Fata et al., 2014; Hill et al., 2014; Sharifi Kia et al., 2020). Effective fiber-ensemble biomechanical properties of combined collagen and myofiber bundles were then obtained for each specimen, using the equibiaxial response (Avazmohammadi et al., 2017b; Hill et al., 2014; Sacks, 2003; Sharifi Kia et al., 2020). This was then used to estimate the effective myofiber and collagen stiffness for each specimen, using a rule of mixtures approach (Avazmohammadi et al., 2017b; Hill et al., 2014; Sharifi Kia et al., 2020):

$$E_{\text{Myofiber}} = \frac{TM_{\text{Before Collagen Recruitment}}}{\Phi_{\text{Myofiber}}} \qquad \text{Eq. 1}$$

$$E_{\text{Collagen}} = \frac{TM_{\text{After Collagen Recruitment}} - (E_{\text{Myofiber}} \times \Phi_{\text{Myofiber}})}{\Phi_{\text{Collagen}}}$$

Here, $E_{\text{Myofiber}}$ and $E_{\text{Collagen}}$ are the effective myofiber and collagen stiffness, respectively, while $TM_{\text{Before Collagen Recruitment}}$ and $TM_{\text{After Collagen Recruitment}}$ are the slope of the lines fitted via linear regression to the fiber-ensemble stress-strain data, before and after collagen recruitment. $\Phi_{\text{Myofiber}}$ and $\Phi_{\text{Collagen}}$ represent the myofiber and collagen area fractions, measured via histological analysis. We assumed the initial portion of the fiber-ensemble stress-strain responses to be mostly dominated by myofibers, while collagen fibers dominated the high-strain response following collagen recruitment (Avazmohammadi et al., 2017a; Hill et al., 2014; Sharifi Kia et al., 2020). As previously described (Fata et al., 2014; Hill et al., 2014), transition strains for categorizing the data before and after collagen recruitment were obtained by differentiating the fiber-ensemble stress-strain response with respect to strain to evaluate the changes in effective fiber-ensemble stiffness. We observed a relatively linear region dominated by myofibers, followed by beginning of collagen recruitment and an abrupt increase in fiber-ensemble stiffness (nonlinear portion of the stress-strain curve) before reaching a plateau at a second linear region (upper limit of collagen recruitment). The data before beginning of collagen recruitment and after the upper limit of recruitment were used to estimate $TM_{Before\ Collagen\ Recruitment}$ and $TM_{After\ Collagen\ Recruitment}$, respectively.

Additionally a nonlinear anisotropic constitutive model (Choi and Vito, 1990) was used to model the response of each specimen:

$$W = B_0(e^{\frac{1}{2}b_1E_{11}^2} + e^{\frac{1}{2}b_2E_{22}^2} + e^{b_3E_{11}E_{22}} - 3) \qquad \text{Eq. 2}$$

Here, W is the strain energy density, $E_{11}$ and $E_{22}$ represent the circumferential and longitudinal (apex to base) Green-Lagrange strains, respectively, $B_0$ is a scaling factor and $b_1$, $b_2$ and $b_3$ are metrics for the circumferential, longitudinal and in-plane coupling stiffness of the RVFW, respectively. 2$^{nd}$ Piola–Kirchhoff stress components were obtained via differentiating equation 2 with respect to Green-Lagrange strain. Specimen-specific model parameters were then estimated based on the experimental stress-strain data, using a trust-region-reflective nonlinear least-squares optimization algorithm in MATLAB (Mathworks, Natick, MA). A $R^2$ measure was used to evaluate the goodness of fit. Age-specific strain energy maps were then generated by taking the average of all strain energy distributions in the circumferential-longitudinal strain space, for RVFW specimens in each age group.

#### 1.3. Quantitative Histological Analysis

Transmural histological staining was performed on a sub-group of specimens from each group ($n_{Control} = 3, n_{Aging} = 3$) to quantify the effects of aging on RV fiber architecture. Based on our previous work showing minimal between-sample variabilities in RV structure (Hill et al., 2014; Sharifi Kia et al., 2020), a sample size of 3 was considered adequate for statistical analysis of our hypothesis. Specimen fixation was carried out using 10% neutral buffered formalin and RVFW specimens were stained using

Masson's trichrome which stains collagen fibers in blue and myofibers in red/pink. A total of 11-17 sections at 50-75 µm increments were obtained for each specimen (from epi to endocardium). Transmural area fractions of collagen and myofibers were then quantified via manual thresholding of the histological images (blue/red) to analyze the effects of aging on RVFW composition. Area fractions were calculated as the ratio of the area occupied by respective blue/red pixels, divided by the total area within the region of interest (ROI). In addition, cardiomyocyte width was measured from the histological data to investigate the role of aging in RV hypertrophy (40 measurements performed on each specimen). Furthermore, for each histological section, local image gradients were used to construct the structure tensor of the gradient map:

$$T = \begin{bmatrix} \iint R(x,y)I_x(x,y)I_x(x,y)dxdy & \iint R(x,y)I_x(x,y)I_y(x,y)dxdy \\ \iint R(x,y)I_x(x,y)I_y(x,y)dxdy & \iint R(x,y)I_y(x,y)I_y(x,y)dxdy \end{bmatrix} \quad \text{Eq. 3}$$

Here, T is the symmetric positive-definite structure tensor, R(x,y) is a gaussian weighing function which specifies the integration ROI and $I_x$ and $I_y$ are the partial spatial derivatives of the histological image (I), respectively, in x and y directions. The 1st eigen vector of T indicates the dominant fiber orientation at each transmural histological section (Hill et al., 2014; Rezakhaniha et al., 2012; Sharifi Kia et al., 2020). For all data presented in this work, 0° corresponds to the circumferential direction, while +90° points towards the apex to base (longitudinal) direction, when looking at the RVFW from the epicardial side. Moreover, collagen fiber coherency was evaluated as:

$$C = \frac{\lambda_1 - \lambda_2}{\lambda_1 + \lambda_2} \times 100 \quad \text{Eq. 4}$$

where $\lambda_1$ and $\lambda_2$ correspond to the 1st and 2nd eigen values of the structure tensor T (Rezakhaniha et al., 2012). 0% collagen fiber coherency corresponds to a sparse (non-coherent), randomly distributed fiber architecture, while 100% coherency indicates a highly-aligned, tightly packed, continuous (coherent) collagen fiber distribution (Clemons et al., 2018).

A total of 66 histological sections were analyzed for the control and aging groups. Transmural data are reported against normalized tissue thickness. We performed linear interpolations to report the histological data on an equally-spaced grid (0-100% thickness). In case of data categorization (Epi, Mid and Endo groups), the data between 0-20% thickness were used for the epicardium, while the data between 80-100% thickness correspond to the endocardium. Orientation analysis and image segmentation were performed using the OrientationJ toolbox (Püspöki et al., 2016; Rezakhaniha et al., 2012) in ImageJ (imagej.nih.gov).
